# Identification and functional characterization of toluene degradation genes in *Acinetobacter* sp. Tol 5

**DOI:** 10.1101/2025.05.12.653411

**Authors:** Shogo Yoshimoto, Maiko Hattori, Shori Inoue, Sakura Mori, Yuki Ohara, Katsutoshi Hori

## Abstract

Microbial degradation of aromatic compounds provides sustainable solutions for environmental remediation and bioconversion. *Acinetobacter* sp. Tol 5 is notable for its strong adhesiveness and potential as a biocatalyst for toluene degradation; however, its toluene metabolic pathway has not been fully elucidated. In this study, genomic analysis identified a cluster of genes in Tol 5 highly similar to the well-known *tod* operon of *Pseudomonas putida*, encoding enzymes responsible for toluene metabolism. Phylogenetic analyses indicated that these *tod* genes, unusual among *Acinetobacter* species, were likely acquired through horizontal gene transfer. Transcriptomic analyses revealed that *todF* and *todC1* are co-transcribed, while the adjacent *fadL2* gene, encoding a putative outer membrane transporter corresponding to *P. putida todX*, is independently transcribed. Functional characterization using gene-knockout mutants revealed that TodC1, the large subunit of dioxygenase, is essential for growth on toluene, whereas FadL2 is not essential. Growth experiments further showed that the *todC1* knockout mutant could metabolize benzoate, but not toluene or benzene, confirming that the TOD pathway is the primary route for toluene and benzene degradation in Tol 5. The identification of the functional TOD pathway, which is unique within *Acinetobacter*, provides genetic and biochemical insights for the development of Tol 5 as an efficient immobilized biocatalyst for the bioremediation and bioconversion of aromatic compounds.

## Introduction

Toluene consumption has been increasing due to its widespread use in industrial applications, including solvents, coatings, adhesives, and fuel additives (1). As a major component of BTEX compounds (benzene, toluene, ethylbenzene, and xylene), toluene is released into air and water, resulting in environmental pollution and raising concerns about toxicity and persistence (2). Given the growing demand for green and cost-effective remediation methods, microbial biodegradation and bioconversion have emerged as sustainable and energy-efficient treatments (3, 4). Unlike chemical treatments, microbial processes can achieve complete degradation of toluene under mild conditions.

Bacteria have evolved diverse metabolic pathways to degrade toluene (5, 6). The well-known pathways involve monooxygenases, which oxidize the aromatic ring or the methyl side chain of toluene, leading to the formation of cresols or benzyl alcohol, which are subsequently converted into catechol-type intermediates by other downstream enzymes. Another well-characterized pathway is the toluene dioxygenase (TOD) pathway. It employs a multicomponent enzyme system that comprises a reductase (*todA*), a ferredoxin (*todB*), and a dioxygenase (*todC1* and *todC2*) that oxidize the aromatic ring of toluene, along with other enzymes (*todD, E, F, G, H*, and *I*) responsible for downstream metabolism (7). This TOD pathway, originally characterized in *Pseudomonas putida* (8), converts toluene to 3-methylcatechol and finally to pyruvate and acetyl-CoA.

While *Acinetobacter* species are not typically recognized as toluene degraders, some strains have been reported to possess toluene-degrading capabilities. For example, *A. junii* isolated from petroleum-contaminated soil degrades 70% of toluene at 0.15 mg/ml within 72 h (9). *A. baylyi*, previously known as *A. calcoaceticus*, completely degrades toluene at 0.3 mg/ml within 14 days (10). *Acinetobacter* sp. Tol 5 isolated from a biofiltration system completely degrades toluene at 0.43 mg/ml within 37 h (11). Among *Acinetobacter*, Tol 5 is unique in terms of its high adhesiveness to various surfaces through its adhesive nanofiber protein AtaA in addition to its high toluene-degradation ability (12, 13). Due to its adhesiveness, Tol 5 cells can be rapidly immobilized onto support and efficiently used for degradation of toluene (14). However, despite the practical potential for biodegradation and bioconversion of toluene, the specific metabolic pathway and genes involved in toluene degradation of Tol 5 remain unclear.

In this study, we elucidated the toluene-degradation pathway of Tol 5. Genomic analysis identified candidate toluene-degrading genes, and gene-knockout mutants verified their function. These findings reveal how Tol 5 metabolizes toluene and provide new insights into toluene metabolism in *Acinetobacter*.

## Materials and Methods

### Strains, plasmids, and culture conditions

The strains and plasmids used in this study are shown in Table S1. *Escherichia coli* and its transformants were cultured in Luria-Bertani (LB) medium (20066-95; Nacalai Tesque) at 37 °C. Tol 5 and its mutants were cultured in LB medium or basal salt (BS) medium (11) at 28 °C.

### Gene knockout

The gene knockout mutants were generated using the cytidine base editing system for *Acinetobacter* according to the previous report (15, 16). The DNA dimer fragment encoding the sequence of the single guide RNA was prepared by mixing oligo DNAs listed in Table S2 in 50 mM NaCl solution and gradually cooling from 95 °C to 18 °C at 0.1 °C/s. This dimer was introduced into the BsaI site of pBECAb-apr. The constructed plasmid was electroporated into Tol 5_REK_, a restriction-modification system and *ataA*-deficient strain (17). After overnight culture of Tol 5_REK_ harboring the plasmid in BS medium to promote mutation, the cells were spread on a BS agar plate containing 5 % (w/v) sucrose to remove the plasmid. Mutation of the target gene was confirmed by sequencing the DNA amplified from the genome by PCR using KOD FX Neo (TOYOBO, Osaka, Japan).

### Bacterial cell growth

The cells cultured in BS medium supplemented with 0.2 % sodium lactate were collected by centrifugation, washed two times with BS medium, and resuspended in BS medium. A 0.25 mL of the suspension was inoculated into 5 mL of BS medium in a glass test tube. The following carbon sources were separately added into each tube: 2.4 μL of toluene or 48 μL of 10 % sodium lactate. The test tubes were sealed with rubber stoppers and incubated at 28 °C with shaking at 200 rpm. After an overnight incubation, photographs of the test tubes were taken to visually assess the bacterial growth.

To make growth curves, the cells cultured in BS medium supplemented with each carbon source were collected by centrifugation, washed two times with BS medium, and resuspended in BS medium. A 0.25 mL of the suspension was inoculated into 5 mL of BS medium in a glass test tube. The following carbon sources were separately added into each tube: 1.2 μL of toluene, 1.0 μL of benzene, or 16 μL of 10 % sodium benzoate, and the optical density was monitored every 60 min using OD-monitorC&T system (TAITEC, Saitama, Japan) during the culture.

### Bioinformatics

To obtain amino acid sequences with similarity to the TodC1 (BCX75806.1) and FadL2 (BCX75810.1), PSI-BLAST searches were performed at an E-value cutoff of 1e-30 against the nr30 or nr90 databases on the MPI Bioinformatics Toolkit (18). The collected sequences were clustered using CLANS (19).

## Results

### Toluene degradation genes of Tol 5

BLAST searches were conducted on the genome sequence of Tol 5 (20) using the toluene degradation genes, *tod*, from *Pseudomonas putida*. The 12 genes with high sequence similarity to the *tod* genes, annotated as *todF, C1, C2, B, A, D, E, G, I, H, S, and T*, were identified. Figure 1 illustrates the genetic organization of the *tod* genes in Tol 5, highlighting its similarity to that of *P. putida*. Surrounding these genes, transposon-related sequences such as a reverse transcriptase, an integrase, and IS elements were observed. When the amino acid sequences of TodC1, the toluene dioxygenase large subunit responsible for the enzyme activity (21), were compared between the two species, they showed a high sequence similarity (identity = 92 %). Other *tod* enzyme genes and the two-component regulatory system TodS and TodT controlling the expression of the *tod* genes (22) also showed high sequence similarities (Table S3). In addition, a gene encoding a putative outer membrane transporter was identified in Tol 5 at the genomic locus corresponding to *todX*, which encodes an outer membrane toluene transporter in *P. putida*. Although this gene was predicted to belong to the outer membrane fatty acid transporter (FadL) family as in *P. putida todX* (23), the sequence similarity with TodX was relatively low (identity = 39 %). Therefore, the gene was designated *fadL2*.

**Fig. 1.**
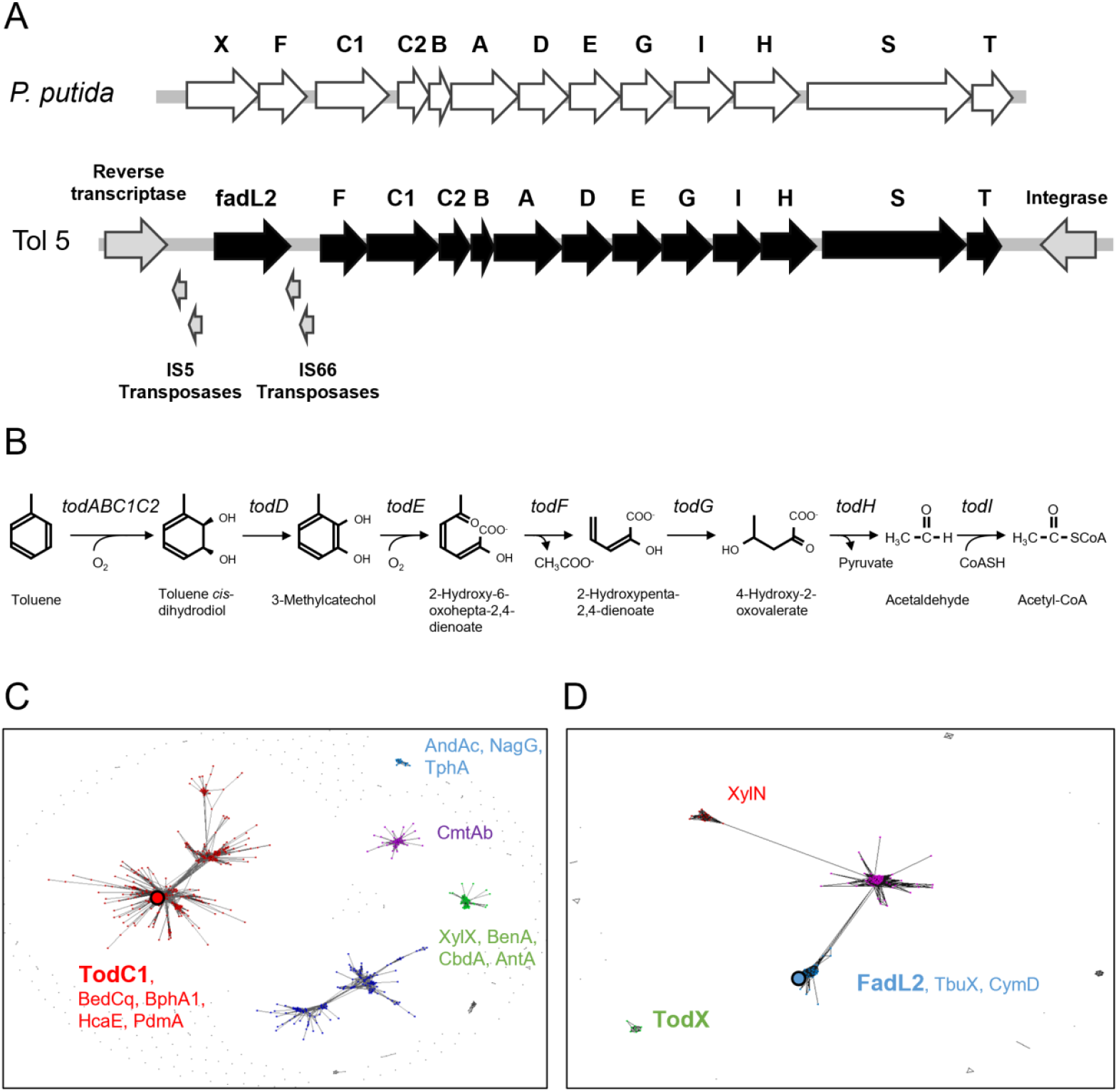
Sequence analysis of the *tod* operon. (A) Genetic map of the *tod* operon region of *P. putida* (top) and *Acinetobacter* sp. Tol 5 (bottom). (B) Reactions catalyzed by the TOD pathway for toluene degradation. (C,D) Clustering analysis of amino acid sequences similar to those of TodC1 and FadL2. Large dots highlight TodC1 and FadL2 of Tol 5. (C) Each dot represents amino acid sequences similar to that of Tol 5 TodC1. (D) Each dot represents amino acid sequences similar to that of Tol 5 FadL2.

To investigate the evolutionary and phylogenetic background of these *tod* genes identified in Tol 5, a BLAST analysis was performed using the amino acid sequence of TodC1 and FadL2 of Tol 5 as a query. Many hits were found in bacteria belonging to the genus *Pseudomonas*, whereas no homologous genes were detected in the genomes of *A. bereziniae*, the closely related species of Tol 5, or *Acinetobacter baumannii*, the most extensively studied species in the genus *Acinetobacter*. Within the current NCBI database, *todC1* and *fadL2* were identified in only three strains of *Acinetobacter*: *A. johnsonii, A. pittii*, and *Acinetobacter* sp. ANC 4635. Although this BLAST analysis highlighted the rarity of the *tod* genes in *Acinetobacter*, it was insufficient to elucidate the evolutionary relationships of *todC1* and *fadL2* from Tol 5 with other aromatic-ring-hydroxylating genes. Therefore, we next conducted a cluster analysis. Amino acid sequences of aromatic ring hydroxylating dioxygenase alpha subunits were collected from the nr30 database using PSI-BLAST and subsequently classified based on their sequence similarity using CLANS, resulting in five distinct clusters (Fig. 1C). While TodC1 of Tol 5 grouped with TodC1 of *P. putida*, biphenyl 2,3-dioxygenase BphA (24), and 3-phenylpropionate dioxygenase HcaE (25), some dioxygenases for toluene were included also in a separate cluster. This seems to reflect the classification of aromatic ring-hydroxylating oxygenases reported in the previous study (26). A similar clustering analysis was performed for FadL2 of Tol 5 (Fig. 1D). FadL2 of Tol 5 was grouped with transporters reported to facilitate the uptake of aromatic compounds, such as TbuX, encoded in the toluene *m*-monooxygenase operon (27), CymD, encoded in the *p*-cumate-2,3-dioxygenase operon (28), and XylN, encoded in the toluene and xylenes monooxygenase operon (29). However, TodX of *P. putida* formed a distinct cluster from these transporters. While the exact genetic origin of *fadL2* remains unclear, it appears to have been derived from a transporter associated with an aromatic oxygenase other than *tod*.

### Gene expression of the *tod* genes in Tol 5

The promoter region of *todX* in *P. putida* has been studied in detail (30). It was shown that the *tod* genes are regulated by the three regions with sequences called TodT-boxes. Comparing this region with the upstream region of *tod* genes of Tol 5, we found that the transcription initiation site is located approximately 200 bases upstream of *todF* of Tol 5, showing high sequence similarity to the *todX* promoter region of *P. putida* (30) (Fig. 2A). Furthermore, the toluene-binding site of TodS predicted in *P. putida* (31) was also conserved in that of Tol 5 (Fig. S1). These results suggest that the expression of the *tod* genes of Tol 5 is regulated similarly by the TodST two-component system, as in *P. putida*. In *P. putida*, the transcription initiation gene is the *todX*, which encodes an outer membrane protein involved in toluene uptake. However, in Tol 5, the presumed transcription initiation gene is *todF*, located downstream of *fadL2* which corresponds to *todX*.

**Fig. 2.**
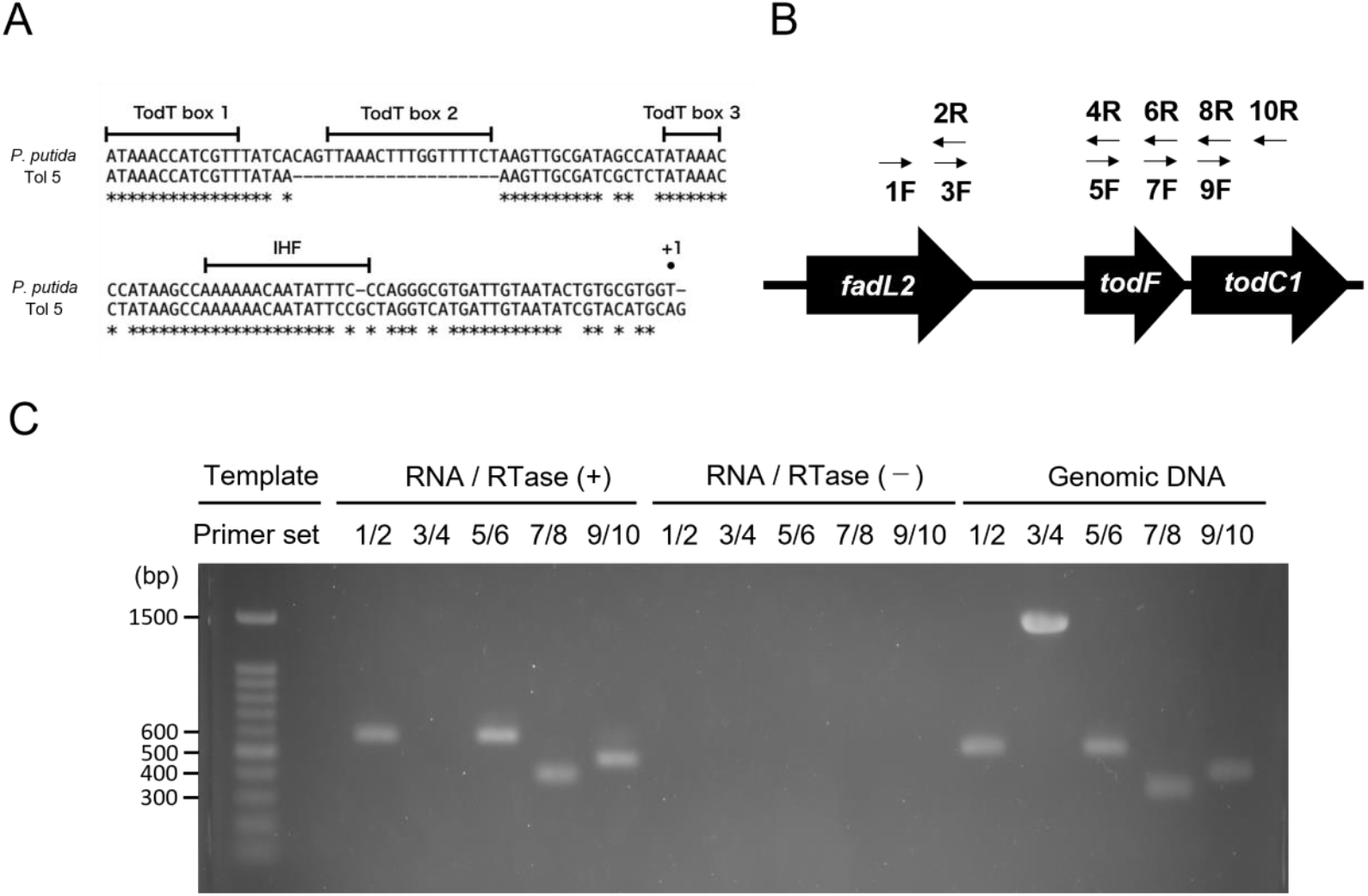
Comparison of promoter regions of *tod* operons. The upstream sequence of *todX* from *P. putida* DOT-T1E was compared with that of *todF* from Tol 5. The ‘+1’ position indicates the transcription initiation site. The integration host factor (IHF) binding site, two pseudopalindromic sequences (box 1 and box 2), and a half-site (box 3), previously identified in *P. putida*, are indicated. (B,C) RT-PCR analysis of total RNA isolated from Tol 5 cells. Primer sets 1F/2R and 5F/6R and 9F/10R were used to examine the transcription of each ORFs. Primer set 3F/4R was used to examine the cotranscription of *fadL2* and *todF*, whereas primer set 7F and 8R was used to examine the cotranscription of *todF* and *todC1*.

To investigate whether *todF* in Tol 5 is transcribed as part of an operon with its neighboring genes, *fadL2* (upstream) and *todC1* (downstream), we performed reverse transcription polymerase chain reaction (RT-PCR) using primer sets targeting regions within and between the three genes. All primer sets successfully amplified DNA fragments from Tol 5 genomic DNA (Fig. 2C; Genomic DNA), confirming the accuracy of the primer design. When using cDNA synthesized from Tol 5 RNA (Fig. 2C; RNA/RTase(+)), we obtained an amplification product for the region between *todF* and *todC1*, suggesting that these genes belong to the same operon. In contrast, no amplification was observed for the region between *todF* and *fadL2*, indicating that these genes are transcribed separately. Furthermore, no PCR amplification was detected in control samples without reverse transcriptase (Fig. 2C; RNA/RTase(−)), demonstrating that the amplification was derived from RNA and not from contaminated genomic DNA. These results suggest that, unlike *P. putida*, the *tod* operon in Tol 5 does not start transcription from *fadL2*, corresponding to *todX*, but instead begins from *todF*.

### Growth abilities of mutant strains on toluene and other carbon sources

To investigate whether the *tod* genes, which are rarely found in other *Acinetobacter* species, are involved in toluene degradation in Tol 5, we constructed knockout mutants of *todC1* and *fadL2* using the Tol 5_REK_ strain (17). Tol 5_REK_ carries mutations in the restriction-modification system, making genetic manipulation easier, and also harbors a mutation in AtaA, resulting in decreased cell adhesion and autoaggregation, thereby improving the accuracy of growth evaluation through turbidity measurements. The growth abilities of Tol 5_REK_ and its Δ*todC1* and Δ*fadL2* mutant strains were compared in basal salt (BS) medium (11). When sodium lactate was added as the sole carbon source to BS medium, all strains grew well. On the other hand, when toluene was added as the sole carbon source, the Δ*todC1* mutant strain failed to grow, while both Tol 5_REK_ and the Δ*fadL2* mutant strain grew well (Fig. 3). These results suggest that *todC1* is involved in toluene degradation, whereas *fadL2* is not essential.

**Fig. 3.**
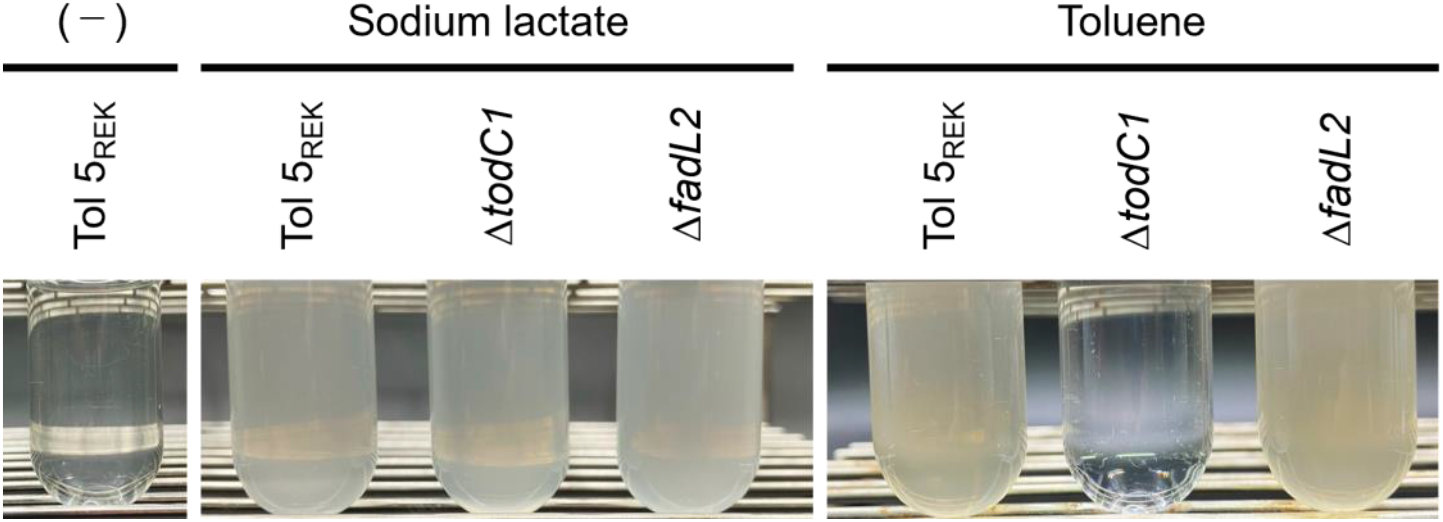
Assimilation test. Overnight culture of Tol 5_REK_, Δ*todC1* mutant, and Δ*fadL2* mutant in BS medium with sodium lactate or toluene. As a control, Overnight culture of Tol 5_REK_ without any carbon source (−) was also shown.

The effect of mutation in *todC1* on the utilization of other carbon sources—benzene, which is metabolized via the TOD pathway, and sodium benzoate, which is metabolized via another pathway, the Ben or Xyl pathway (32, 33)—was evaluated by measuring cell growth. In BS medium containing toluene or benzene as the sole carbon source, the Δ*todC1* mutant strain showed no growth (Fig. 4). On the other hand, when sodium benzoate was added as the sole carbon source, both Tol 5_REK_ and the Δ*todC1* mutant strain grew well. These results indicate that *todC1* plays a crucial role in the degradation of toluene and benzene but is not involved in the utilization of benzoate and lactate. This also suggests that the TOD pathway is the primary route for the degradation of both toluene and benzene in Tol 5.

**Fig. 4.**
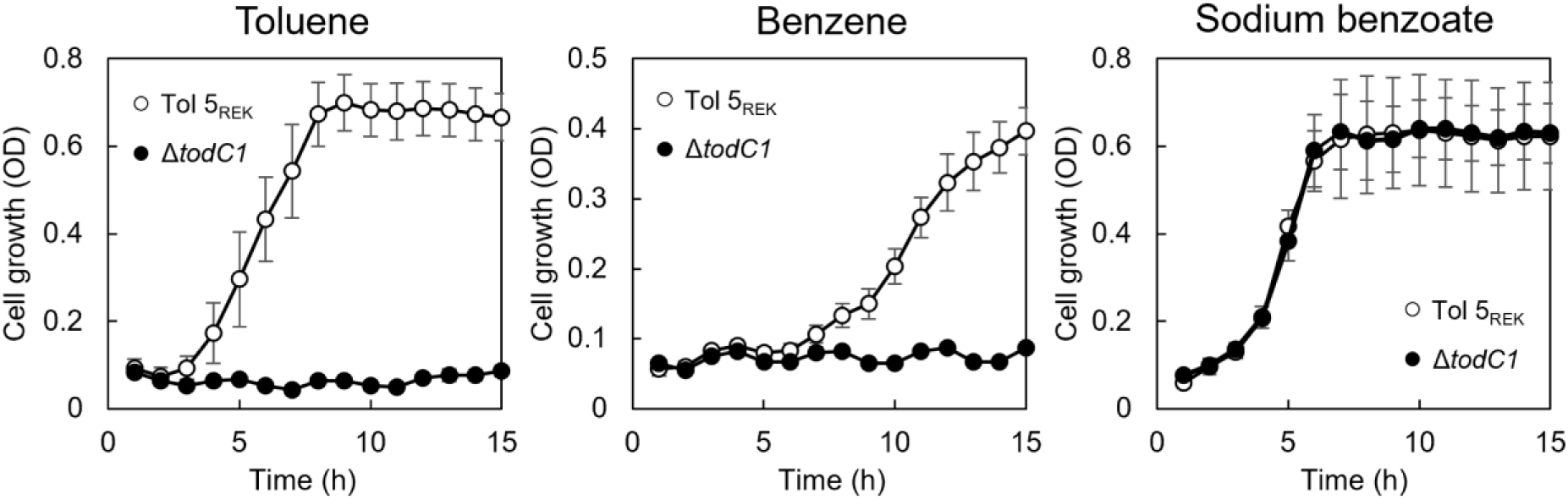
Growth curves with different carbon sources. Tol 5_REK_ (open circle) and the Δ*todC1* mutant strain (closed circle) was inoculated to BS medium containing toluene, benzene, or sodium benzoate. The data are presented as the means ± SEMs (n = 3).

## Discussion

In this study, we discovered that Tol 5 possesses a gene cluster for toluene degradation highly similar to the *tod* operon of *P. putida* (Fig. 1). Genomic analysis revealed that this gene island encodes homologs of the TOD pathway enzymes (*todF, C1, C2, B, A, D, E, G, I, H*) and the two-component regulatory system (*todS* and *todT*). Our experimental data showed that the Δ*todC1* mutant strain lost the ability to grow on toluene and benzene (Fig. 3 and 4), confirming that the TOD pathway is the primary route for Tol 5 to degrade those aromatic hydrocarbons. This is the first report of a functional *Pseudomonas*-like TOD pathway identified in *Acinetobacter*.

Bioinformatic analysis suggests that Tol 5 acquired its *tod* genes through horizontal gene transfer. Transposon-like sequences identified at the boundary of the *tod* genes (Fig. 1A) indicate transposition events facilitating their integration. Furthermore, BLAST searches show that these *tod* genes are uncommon among *Acinetobacter* species, supporting the hypothesis that the *tod* genes in Tol 5 were horizontally acquired. Indeed, horizontal gene transfer is recognized as a key mechanism by which bacteria acquire gene clusters involved in the degradation of organic pollutants (34). Horizontal transfer of such a large metabolic operon could immediately confer new metabolic capabilities, explaining how Tol 5 came to degrade toluene effectively.

In *P. putida, todX*, encoding an outer membrane protein thought to facilitate toluene uptake, is co-transcribed as part of the *tod* operon (23). The Tol 5 genome encodes *fadL2*, a putative outer membrane transporter, in place of *todX*. However, the sequence similarity of FadL2 with TodX is relatively low (Table S3) and *fadL2* is transcribed independently of the *tod* operon (Fig.2). In addition, unlike *todC1, fadL2* was not essential for toluene degradation (Fig. 3 and 4). These findings suggest that Tol 5 may employ an alternative mechanism for toluene uptake, possibly involving an unidentified transporter or passive diffusion across the membrane.

The strong adhesion ability of Tol 5, due to AtaA, has been utilized for rapid immobilization in bioreactors (14, 35). Now, with the identification of the *tod* pathway, Tol 5 could be a promising biocatalyst for transforming toluene into valuable compounds. Further genetic modifications, such as disruption of *tod* genes, could enable the production of industrially relevant intermediates (36). The combination of such genetic modifications and cell immobilization techniques could lead to the development of efficient bioprocesses using Tol 5 as an immobilized biocatalyst.

In conclusion, our findings provide a detailed genetic and functional characterization of the *tod* operon in Tol 5, revealing its potential for industrial applications such as bioremediation and the bioconversion of toluene into high-value chemicals.

## Acknowledgements

The authors thank Michio Homma for helpful discussion. The authors also thank Eriko Kawamoto for her technical assistance. This research was supported by the Graduate Program of Transformative Chem-Bio Research at Nagoya University supported by MEXT (WISE Program) to SI, the Japan Society for the Promotion of Science (JSPS) KAKENHI (Grant Number JP24H00043) to KH and SY, and GteX Program Japan Grant number JPMJGX23B4 to KH.

