## Supporting Information for "Identification and functional characterization of toluene degradation genes in *Acinetobacter* sp. Tol 5"

Short title: Toluene degradation genes in *Acinetobacter* sp. Tol 5

Shogo Yoshimoto[1], Maiko Hattori[1], Shori Inoue[1], Sakura Mori[1], Yuki Ohara[2], Katsutoshi Hori[1,*]

[1]Department of Biomolecular Engineering, Graduate School of Engineering, Nagoya University, Chikusa-ku, Nagoya, Aichi 464-8603, Japan.

[2]Friend Microbe Inc. Chikusa-ku, Nagoya, Aichi 464-0858, Japan.

```
Tol 5       MSFWDRRKPQNKLKNNHYNISLKEKGSAELMQEQYTRIIFDGLYEFVGLLDAKGNVLEVN
P.putida    MSSLDRKKPQNRSKNNYYNICLKEKGSEELTCEEHARIIFDGLYEFVGLLDAHGNVLEVN
            **  ** ****  *** *** ****** **  *   **************** *******

Tol 5       QVALEGAGITLEDIQGKPFWKTPWWQLSKKTMATQKRLIETASSGEFVRCDVEILGKADG
P.putida    QVALEGGGITLEEIRGKPFWKARWWQISKKTEATQKRLVETASSGEFVRCDVEILGKSGG
            ****** ***** * ******  *** **** ****** ******************  *

Tol 5       KEIIAVDFSLLPIRNEEGEIIYLLAEGRNITDKKKAEAMLALKNQELEQSVEYIRKLDKQ
P.putida    REVIAVDFSLLPICNEEGSIVYLLAEGRNITDKKKAEAMLALKNQELEQSVECIRKLDNA
             * ********** **** * ******************************* *****
```

**Fig. S1.** Sequence alignment of TodS from *Acinetobacter* sp. Tol 5 and *P. putida*. The toluene-binding residues in TodS from *P. putida* were highlighted in yellow.

Table S1. Strains and plasmids used in this study.

| Strain | Description | Reference |
|---|---|---|
| *Acinetobacter* sp. | | |
| Tol 5 | Wild type strain | 1 |
| Tol 5 $_{REK}$ | Restriction enzyme-encoding genes and *ataA* knockout mutant of Tol 5, REK123Δ*ataA* | 2 |
| Tol 5 $_{REK}$ Δ*todC1* | *todC1* gene knockout mutant of Tol 5$_{REK}$ | This study |
| Tol 5 $_{REK}$ Δ*fadL2* | *fadL2* gene knockout mutant of Tol 5$_{REK}$ | This study |
| *Escherichia coli* | | |
| DH5α | Host for routine cloning | Purchased from Takara (Shiga, Japan) |
| Plasmid | | |
| pBECAb-apr | For gene knockout | 3 |
| pBECAb-apr-todC1 | For *todC1* gene knockout | This study |
| pBECAb-apr-fadL2 | For *fadL2* gene knockout | This study |

Table S2. Oligo DNAs used in this study.

| Name | Sequence (5’ to 3’) |
|---|---|
| TodC1-spacer-F | tagtCTGAACCAGTGCCGCCATCG |
| TodC1-spacer-R | aaacCGATGGCGGCACTGGTTCAG |
| FadL2-spacer-F | tagtATCTCAGGGTATGGGAGGAA |
| FadL2-spacer-R | aaacTTCCTCCCATACCCTGAGAT |

Table S3. Amino acid similarities of *tod* genes from Tol 5 and *P. putida*.

| Protein | Identity (%) |
|---|---|
| TodX / FadL2 | 39 |
| TodF | 92 |
| TodC1 | 92 |
| TodC2 | 90 |
| TodB | 93 |
| TodA | 84 |
| TodD | 93 |
| TodE | 81 |
| TodG | 86 |
| TodI | 91 |
| TodH | 97 |
| TodS | 86 |
| TodT | 88 |
